# The DEAD box RNA helicase Ddx39a is essential for myocyte and lens development in zebrafish

**DOI:** 10.1101/209650

**Authors:** Linlin Zhang, Beibei Li, Yuxi Yang, Ian C. Scott, Xin Lou

**Affiliations:** Model Animal Research Center, Nanjing University, China.; Program in Developmental and Stem Cell Biology, The Hospital for Sick Children; Department of Molecular Genetics, University of Toronto, Canada.

**Keywords:** RNA helicase, myocyte, lens, RNA splicing, zebrafish

## Abstract

RNA helicases from the DEAD-box family are found in almost all organisms and have important roles in RNA metabolism including RNA synthesis, processing and degradation. The function and mechanism of action of most of these helicases in animal development and human disease are largely unexplored. In a zebrafish mutagenesis screen to identify genes essential for heart development we identified a zebrafish mutant, which disrupts the gene encoding the RNA helicase DEAD-box 39a (*ddx39a*).Homozygous *ddx39a* mutant embryos exhibit profound cardiac and trunk muscle dystrophy, along with lens abnormalities caused by abrupt terminal differentiation of cardiomyocyte, myoblast and lens fiber cells. Further investigation indicated that loss of *ddx39a* hindered mRNA splicing of members of the *kmt2* gene family, leading to mis-regulation of structural gene expression in cardiomyocyte, myoblast and lens fiber cells. Taken together, these results show that Ddx39a plays an essential role in establishment of proper epigenetic status during cell differentiation.

## Introduction

The DEAD-box RNA helicase family is a large protein group characterized by the presence of an Asp-Glu-Ala-Asp (DEAD) motif that is highly conserved from bacteria to man (Bleichert and Baserga, 2007; Rocak and Linder, 2004). Using the energy derived from ATP hydrolysis, these proteins modulate RNA topology and association/dissociation of RNA-protein complexes. DEAD-box RNA helicases play important roles in all aspects of RNA metabolism, including transcription, pre-mRNA splicing, rRNA biogenesis, RNA transport and translation (Calo et al., 2015; Jarmoskaite and Russell, 2011, 2014; Linder and Jankowsky, 2011). Recently, a fuller appreciation of the functions of these RNA helicases in variant physiological or developmental scenarios has emerged. Numerous reports have described deregulation of expression or function of DEAD-box RNA helicases in cancer development or progression, indicating that DEAD box proteins may be involved in processes that are key to cellular proliferation (Fuller-Pace, 2013; Sarkar and Ghosh, 2016). Recent reports have further shown that DEAD-box RNA helicases play diverse roles in developmental events ranging from body axis establishment (Meignin and Davis, 2008) to germ cell, blood, digestive organ and brain development (Hirabayashi et al., 2013; Hozumi et al., 2012; Payne et al., 2011; Zhang et al., 2012). These studies indicated that DEAD-box RNA helicase family members could regulate specific developmental processes by affecting pre-mRNA splicing, ribosomal biogenesis or RNA transport.

As a member of DEAD box family of ATP-dependent RNA helicases, Ddx39a was originally identified in a screen for proteins that interact with the essential splicing factor U2AF65 (Fleckner et al., 1997). Studies in different organisms indicated that besides playing an important role in pre-mRNA splicing (Fleckner et al., 1997; Shen et al., 2008; Shen et al., 2007), DDX39a also acts in mRNA nuclear export (Luo et al.,2001), cytoplasmic mRNA localization (Meignin and Davis, 2008) and maintenance of genome integrity (Yoo and Chung, 2011). However, how Ddx39a functions during embryogenesis, and which RNAs are processed by Ddx39a in different development scenarios remains open to investigation.

In the current study, we examined the function of *ddx39a* in vertebrate development by using a newly identified zebrafish *ddx39a* gene trap line. Transcriptome profiling and phenotypic analysis of the *ddx39a* mutant showed that *ddx39a* is indispensable for the development of heart, trunk muscle and eyes. Further experiments revealed that Ddx39a can bind to mRNAs encoding a set of epigenetic regulatory factors, including members of KMT2 family. Loss of *ddx39a* leads to aberrant pre-mRNA splicing of these epigenetic regulator transcripts, leading to a failure in establishment of proper epigenetic status of multiple structural genes, and eventually hampers the terminal differentiation of cardiomyocyte, myoblast and lens fiber cell lineages.

## Materials and methods

### Zebrafish lines

Zebrafish embryos were maintained and staged using standard techniques (Westerfield, 1993). The RP-T gene-trap vector was modified from RP2 (Clark et al., 2011). RP-T plasmid (25 ng/μl) and Tol2 transposase mRNA (50 ng/μl) were injected (1 nl each) into 1-cell embryos as previous described (Hou et al., 2017). To identify the affected gene in the gene trap lines, inverse PCR and 5’RACE were applied as previously described (Clark et al., 2011).

### Immunocytochemistry

Embryos were fixed in 4% paraformaldehyde (PFA) at 4 °C overnight. Embryos were blocked with blocking solution (1× PBS, 1% BSA, 1% Triton-X100, 0.1% DMSO) for 2 hours then incubated with primary antibodies diluted in block solution overnight at 4 °C. Embryos were washed 3 times in 1× PBS with 1% Triton-X100 for 15 min. Embryos were incubated with secondary antibodies diluted in blocking solution for 2 hours. Primary antibodies specific to MYH1E (MF-20, obtained from DHSB, 1:200 dilution), F-actin (Abcam, 1:1000 dilution), and Acetylated Tubulin (Sigma, 1:1000 dilution) were used. Fluorescent immunocytochemistry was performed using anti-mouse antibody conjugated with Cy3 (1:1000 dilution, Sigma). Unless otherwise stated, manipulations were performed at room temperature.

### Imaging

Imaging was performed using a Leica DFC320 camera on a Leica M205FA stereomicroscope. All confocal images were taken using a Zeiss LSM880 confocal microscope.

### mRNA injections

pCS2+ vectors carrying a cDNA fragment encoding membrane RFP, Ddx39a, Flag-Ddx39a and Tol2-transposase were used in this study. Capped mRNA was synthesized using an SP6 mMESSAGE mMACHINE kit (Ambion, Life Technologies Corp.). For phenotypic rescue experiments, *ddx39a* mRNA (100 pg) was injected at the one-cell stage.

### RNA *in situ* hybridization

Transcription of DIG-labeled antisense RNA probes was performed using standard methods. RNA *in situ* hybridization (ISH) was carried out as previously described (Thisse and Thisse, 2008).

### RNA-Seq for Expression and Splicing Analysis

Total RNA from 100 control and *ddx39a* mutant 24 hpf embryos was isolated by TRIzol reagent (Invitrogen). Two biological replicates for each group (control and *ddx39a* mutant) were processed and sequenced. Sequencing libraries were prepared using the Nextera sample preparation kit (Illumina) and subjected to HiSEQ paired-end 100 bp plus sequencing. Resulting reads were aligned to the zebrafish reference genome (GRCz10) and gene expression quantified using tophat V2.2.1 and bowtie2 v2.2.3 (Kim et al., 2013; Langmead and Salzberg, 2012). Gene differential expression was analyzed using HTSeq v 0.6.1 p1 (Anders et al., 2015). Genes showing altered expression with adjusted P < 0.05 were considered differentially expressed. For the set of differentially expressed genes a functional analysis was performed using Ingenuity Pathway Analysis Software and DAVID (Huang da et al., 2009), and some of the enriched processes were selected according to relevant criteria related to the biological process studied. Using a R visualization package called GOPlot (Walter et al., 2015), a chord plot was generated to better visualize the relationships between genes and the selected enriched processes. OLego (Wu et al., 2013) and Quantas pipelines were used for alternative splicing analysis. Transcript structure was inferred between paired-end reads. Alternative splicing (AS) was quantified by separating genomic and junction reads and scoring the output from transcript inference. Finally, statistical tests were run to filter the significant AS events [Fisher’s exact test and Benjamini false discovery rate (FDR)]. RNA-seq data have been uploaded to the Gene Expression Omnibus (GEO: GSE97067).

### RNA immunoprecipitation and RIP-seq

RNA immunoprecipitation (RIP) was performed as previously described (Jain et al.,2011) with the specific modifications below. Ddx39a-Flag mRNA was injected into embryos from *ddx39a* heterozygous fish in-crosses, with mutant embryos sorted based on GFP brightness. De-yolked embryos were homogenized in RIP buffer and briefly sonicated using a probe-tip Branson sonicator to solubilize chromatin. Each sample was normalized for total protein amount then Flag-Ddx39a and associated RNA was isolated via incubation with anti-Flag agarose beads (Sigma) for 6 hours at 4 °C with gentle rotation. Samples were washed sequentially in high stringency buffer, high salt buffer and RIP buffer. Ddx39a-associated RNA was extracted with Trizol and then processed for sequencing. Sequencing libraries were prepared using the Nextera sample preparation kit (Illumina) and subjected to HiSEQ paired-end 100 bp plus sequencing. Data analysis was performed as previously described.

### Quantitative real-time PCR

Total RNA was prepared using TRIzol (Invitrogen, Life Technologies Corp.), with DNase-treated RNA reverse transcribed using random 16-mer priming and SuperSript II reverse transcriptase (Life Technologies Corp.). Quantitative PCR (qPCR) assays were performed in triplicate using the SYBR Green Master Mix (Takara) according to the manufacturer’s instructions. The amplified signals were confirmed to be a single band by gel electrophoresis, and they were normalized to the signals of zebrafish 18S rRNA. Primers used for qPCR analysis are listed in Supplementary Table 1.

### Reverse transcription-PCR analysis of splicing

Total RNA was prepared from control and *ddx39a* mutant larvae at 36 hpf. Reverse transcription (RT)-PCR was performed using total RNA prepared as described above to monitor the splicing of pre-mRNAs. The primer pairs and detailed PCR conditions used to amplify each of the genes are listed in Supplementary Tables 1 and 2.

### ChIP-qPCR

ChIP assays were performed as previously described (Lindeman et al., 2009). In brief, at 24hpf, *ddx39a* mutant and control embryos were collected, de-yolked and cross-linked with 1% formaldehyde for 10 min at room temperature and subsequently quenched with glycine to a final concentration of 0.125 M for another 10 min. Chromatin was sonicated with a Bioruptor (Diagenode), cleared by centrifugation, and incubated overnight at 4 °C with 5 mg anti-H3K4Me3 antibody (Abcam). Immunocomplexes were immobilized with 100 ul of protein-G Dynal magnetic beads (Abcam) for 4 hours at 4 °C, followed by stringent washes and elution. Eluates were reverse cross-linked overnight at 65 °C and deproteinated. DNA was extracted with phenol chloroform, followed by ethanol precipitation. H3K4me occupied regions at 24 hfp was retrieved from previously reported data set(Bogdanovic et al., 2012). ChIP-qPCR analyses were performed using a Light Cycler 480II machine (Roche). ChIP-qPCR signals were calculated as percentage of input. All primers used in qPCR analyses are shown in Table S1.

## Results

### Loss of ddx39a leads to an embryonic lethal phenotype in zebrafish

In a *Tol2* transposon-mediated gene-trapping screen to identify novel genes involved in cardiovascular system development (Hou et al., 2017), we identified a zebrafish line, RT-011, in which embryos demonstrated a dynamic GFP expression pattern, with strong signal evident in somites and eyes from the 8 somite stage. As development proceeded, GFP signal also emerged in heart (Fig. 1A). Multiple incrosses of RT-011 heterozygotes yielded wildtype (WT), heterozygous, and homozygous mutant embryos at Mendelian ratios, with all homozygous mutant embryos dying by 4 days post-fertilization (dpf). This indicated that the RT-011 trap line carried a recessive lethal allele. Homozygous mutant embryos from incrosses of heterozygous RT-011 fish showed no obvious morphological defects until after 24 hours post-fertilization (hpf) (Fig. 1B), at which point contraction of the definitive heart tube was extremely weak and irregular in mutants (Movie S1). Homozygous mutant embryos were completely paralyzed at 24 hpf, and did not exhibit spontaneous tail movements or response to tactile stimulation (Movie S2). At later stages of development further defects became apparent, including a failure to establish blood circulation, a curved body axis, a prominent cardiac edema and disorganized myotome (Fig. 1B). 5’ RACE was used to identify the gene trapped in the RT-011 line. Sequencing indicated that the gene-trapping element was integrated within the 2nd intron of *ddx39a* locus (Fig. 1C). The zebrafish Ddx39a protein contains DEXDc (for ATP binding and hydrolysis) and HELICc (helicase superfamily c-terminal) domains; both of which are highly conserved (over 90% amino acid identity) between zebrafish and human (Fig. S1). The gene trap insertion resulted in a transcript that encoded a fusion protein containing the first N-terminal 69 amino acids of Ddx39a (Fig. 1D). Since this fusion protein lacked both the DEXDc and HELICc domains, the allele we identified from the RT-011 line should act as a true null allele. To confirm that mutation of *ddx39a* represents the causal event in the homozygous RT-011 phenotype, mRNA encoding wild-type Ddx39a was injected into embryos from incrosses of heterozygous RT-011 fish. Morphological homozygous RT-011 defects were efficiently rescued by *ddx39a* RNA (Fig. 1E), with injected mutant embryos surviving up to 9 dpf. Taken together, this indicated that developmental abnormalities in RT-011 homozygous embryo resulted from mutation of *ddx39a*.

**Figure 1.**
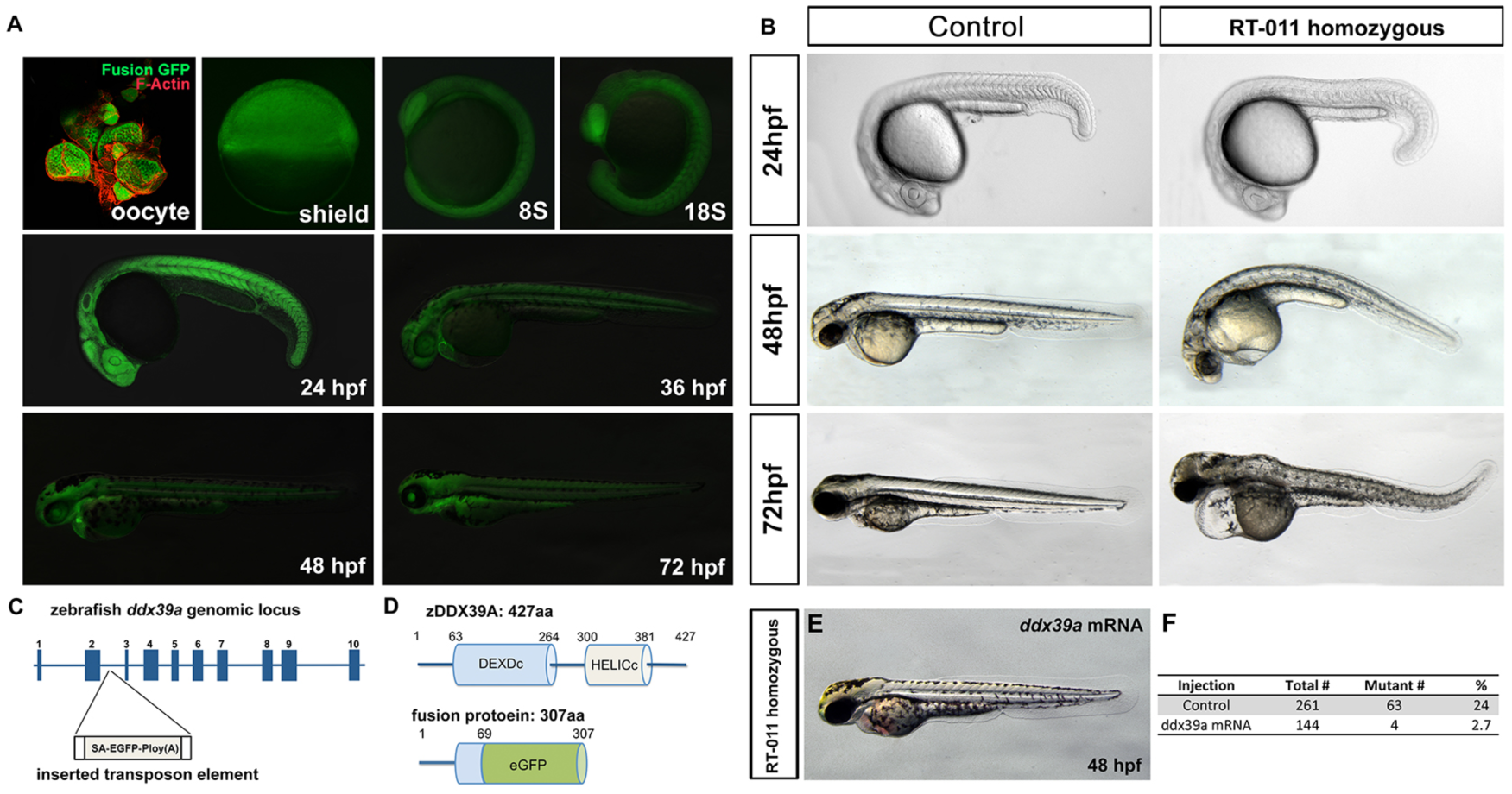
Loss of *ddx39a* leads to an embryonic lethal phenotype in zebrafish. (A): GFP expression pattern in zebrafish trapping line RP24-011. Lateral views, hpf, hour past fertilization. (B): Morphological defects in RP24-011 homozygous mutant embryos: RP24-011 homozygous embryo did not show morphological defect till 24 phf; at 48 hpf, RP24-011 homozygous embryo showed curved body axis; at 72 hpf, RP24-011 homozygous embryo showed heart edema, dysmorphic jaw and global degeneration. (C): The zebrafish *ddx39a* genomic locus. The transposon is inserted after the second intron. (D): Zebrafish DDX39A protein contains a DEXDc domain and a HELICc domain. The insertion of gene trapping element resulted a fusion transcript which will be translated into a fusion protein contain first 69 amino acid from fish DDX39A on the N terminal. (E): Injection of *ddx39a* mRNA rescue the developmental defects in RP24-011 homozygous embryo. (F): The counting of the percentage of embryo showed development defects in a clutch.

To further confirm the endogenous embryonic expression of *ddx39a*, whole mount RNA *in situ* hybridization was carried out on wild type embryos. *Ddx39a* demonstrated strong expression during early embryogenesis, with abundant transcript localized to the myotome, heart and eyes. As development proceeded, the expression of *ddx39a* was constrained to several tissues including pharyngeal arches and liver (Fig. S2). The dynamic expression pattern of *ddx39a* indicated possible roles in multiple developmental stages and tissues.

### Developmental defects in muscular organs and lens in ddx39a mutants

Based on the expression pattern during embryogenesis and defects displayed in *ddx39a* homozygous mutants, we examined development of the heart, skeletal muscle and lens in *ddx39a* mutant embryos.

At 36 hpf, the heart tube in wild type embryos had undergone looping and chamber ballooning, becoming a functional two-chamber pump. In contrast, hearts of *ddx39a* mutants were morphologically abnormal, with little looping or chamber emergence evident (Fig. 2A). Gene expression analysis of a number of cardiac markers at 26 hpf demonstrated that loss of *ddx39a* did not cause a readily apparent decrease in expression of many cardiogenic regulatory genes, including *nkx2.5*, *gata5*, *tbx5a* and *bmp4* (Fig. 2B and data not shown. Specification of both heart chambers occurred properly in *ddx39a* mutants, as shown by expression of *myh6* and *myh7* (Fig. 2C). In contrast, *ddx39a* mutant embryos demonstrated significantly reduced expression of *nppa*, a gene associated with maturation of the heart tube (Auman et al., 2007). Expression of a number of cardiac sarcomere structural genes was also distinctly down-regulated in *ddx39a* mutant embryos, including *myh6*, *cmlcl* and *actalb* (Fig. 2C). Loss of these cardiac sarcomere components may represent the causal factor for the weak contractility observed in *ddx39a* mutant hearts.

**Figure 2.**
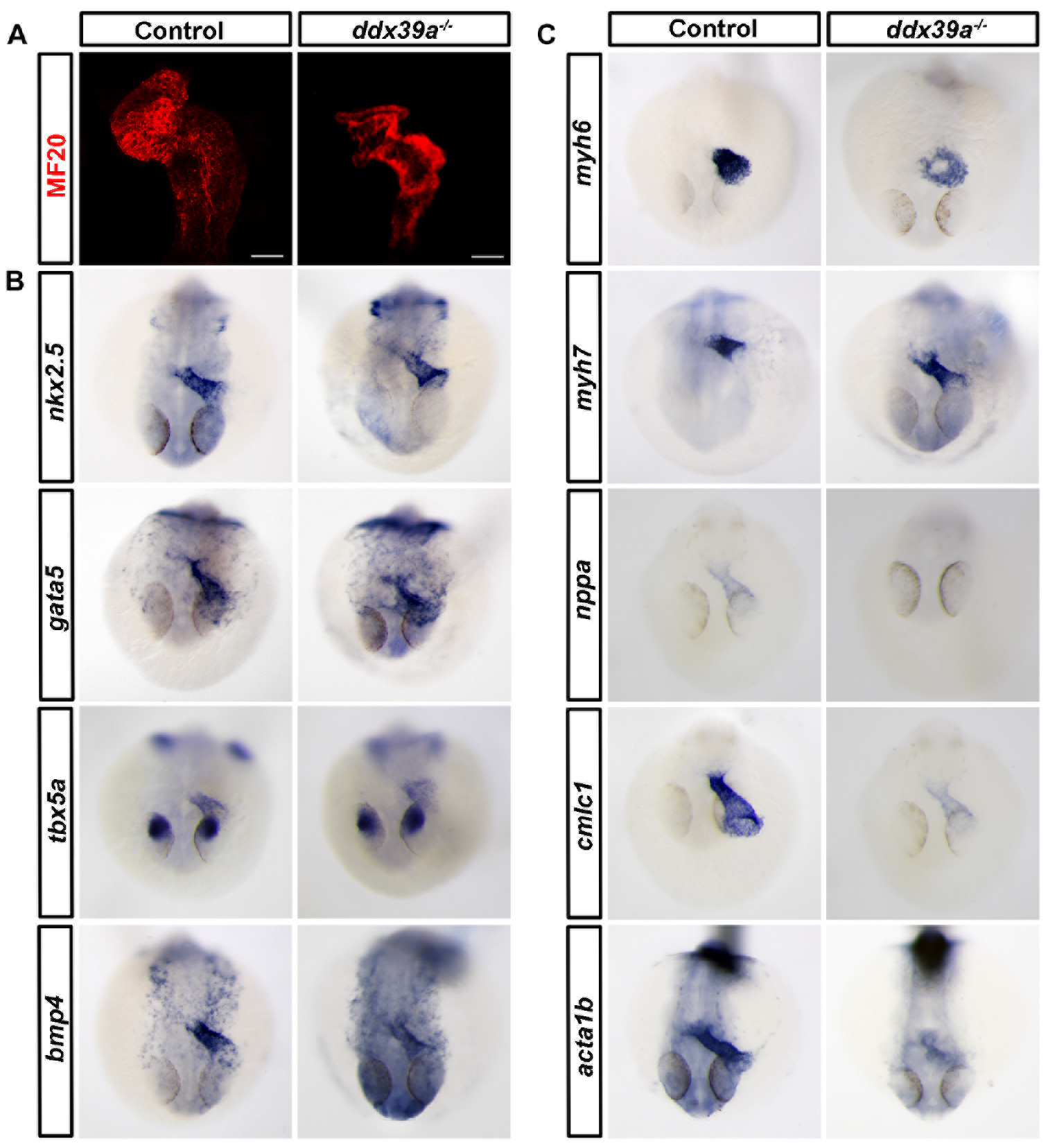
Loss of *ddx39a* led to cardiomyocyte differentiation defects in the zebrafish embryos. (A): Ventral view of heart from whole-mount wide type or *ddx39a* mutant zebrafish embryos at 36 hpf, with cranial to the top. Myocardium was labeled with MF20 antibody. Scale bars: 200um. (B): RNA *in situ* hybridization is shown for cardiogenic regulatory gene expression in WT and *ddx39a* mutant zebrafish embryos. (C): RNA *in situ* hybridization is shown for cardiomyocyte structural gene expression in WT and *ddx39a* mutant zebrafish embryos. B and C, embryos were collected at 26 hpf, frontal views with dorsal side to the top.

Next, we determined whether the locomotion defect in *ddx39a* mutant embryos reflected altered skeletal muscle organization. Immunohistochemical staining for Myosin Heavy Chain (MF20 antibody) and F-Actin revealed myofibrillar protein assembly was severe disrupted in *ddx39a* mutant embryos when compared with wild type (Fig. 3A). To analyze which step of muscle development was affected, *ddx39a* mutant embryos were analyzed for myoblast differentiation via RNA *in situ* hybridization for expression of genes encoding myogenic regulatory factors (Bentzinger et al., 2012). Expression of early myogenic specification markers (*myoD* and *myf5*) and late differentiation markers (*myogenein* and *myf6*) appeared normal until 32 hpf (Fig. 3B and data not shown).

**Figure 3.**
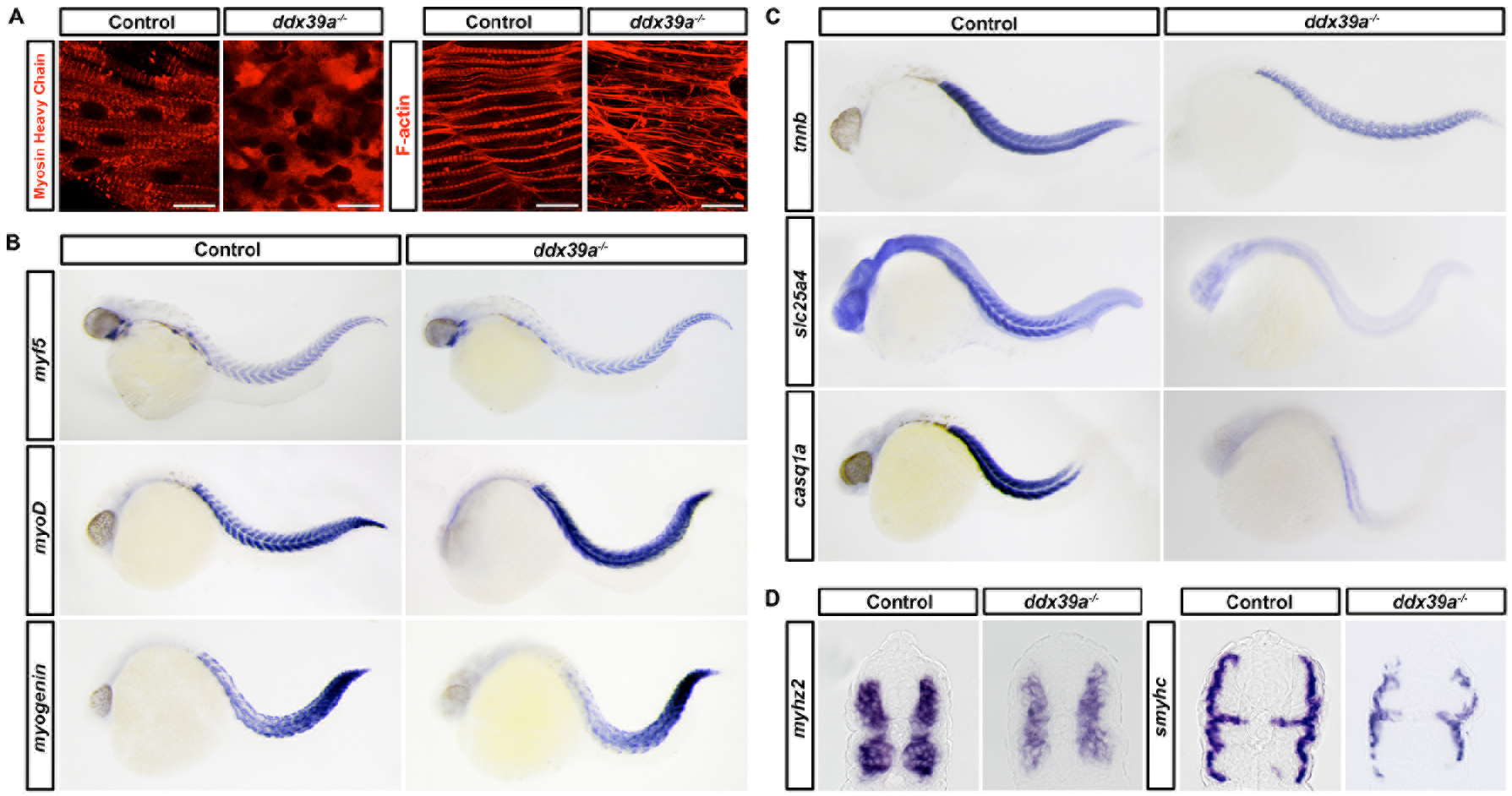
Loss of *ddx39a* resulted in defected skeletal muscles differentiation in zebrafish embryos. (A): Immunostaining showed the organization of myofilaments in wild type or *ddx39a* mutant embryos at 24 hpf. Anti-MHC antibody (MF20) labeled thick (myosin) filaments and anti-actin antibody labeled thin filaments. Scale bars: 20um. (B): RNA *in situ* hybridization is shown for myogenic regulatory gene expression in WT and *ddx39a* mutant zebrafish embryos. (C and D): RNA *in situ* hybridization is shown for myocyte structural gene expression in WT and *ddx39a* mutant zebrafish embryos. B, C and D, embryos were collected at 32 hpf. B and C, lateral views with anterior to the left. D, cross section with dorsal side to top.

Similar to the myocardium, RNA *in situ* revealed that in *ddx39a* mutant embryos the expression of a battery of sarcomeric components, including *tnnb*, *myhz2* and *smyhc,* were significantly reduced (Fig. 3 C and D). This indicated that maturation of both fast muscle and slow muscle was compromised in *ddx39a* mutants. We also found reduction in transcript levels of *casq1a*, which encodes a calcium-binding protein of the skeletal muscle sarcoplasmic reticulum (Yazaki et al., 1990), and *slc25a4*, encoding a muscle cell specific mitochondrial ATP-ADP carrier (Gutierrez-Aguilar and Baines, 2013), in *ddx39a* mutant embryos (Fig. 3 C). These results suggested that *ddx39a* plays an important role in skeletal muscle maturation and function.

As *ddx39a* demonstrated strong expression in developing eyes, we next examined development of the lens in *ddx39a* mutants. At 24 hpf, the lens of *ddx39a* mutant embryos displayed no obvious defects based at the level of gross morphology (Fig. 4A). During eye morphogenesis, cells in the center of the lens mass retain spheroidal morphology; differentiating as primary fibers (which originate from the central lens placode) that elongate and wrap around the ovoid-shaped cells in the center, resulting in a crescent shaped layers of fibers (Greiling and Clark, 2012). In contrast to wild type, in *ddx39a* mutants a majority of lens fiber cells showed relatively irregular shape and did not form crescent shaped layers (Fig. 4A, lower panels). We next analyzed expression of lens-specific genes to explore the nature of the disorganization of primary fiber cells in *ddx39a* mutant embryos. We first examined the expression of a cascade of upstream transcription factors that drive fiber cell differentiation, including *prox1a*, *foxe3* and *pitx3* (Cvekl and Duncan, 2007; Greiling and Clark, 2012; Pillai-Kastoori et al., 2015). Results from *in situ* hybridization showed that in *ddx39a* mutants the transcription of *foxe3* was mildly upregulated, whereas no overt change in *pitx3* and *prox1a* expression were observed (Fig. 4B). During lens fiber cell elongation, soluble proteins known as crystallins are abundantly expressed in lens fibers to increase the refractive index and contribute to transparency (Clark, 2004). We found expression of crystallin genes, including *cryaa*, *cry2d10* and *cryg2d1*, was dramatically down-regulated in *ddx39a* mutant embryos (Fig. 4C). Differentiated lens fiber cells express a set of cellcell adhesion molecules required to create refractive index matching of lens membranes and cytoplasm (Bassnett et al., 2011). *In situ* hybridization demonstrated that in *ddx39a* mutant embryos the expression of *lim2.4*, a lens specific receptor for *calmodulin* involved in cell junction organization, was notably down-regulated (Fig. 4C). These data indicated *ddx39a* is also indispensable for lens fiber cell terminal differentiation.

**Figure 4.**
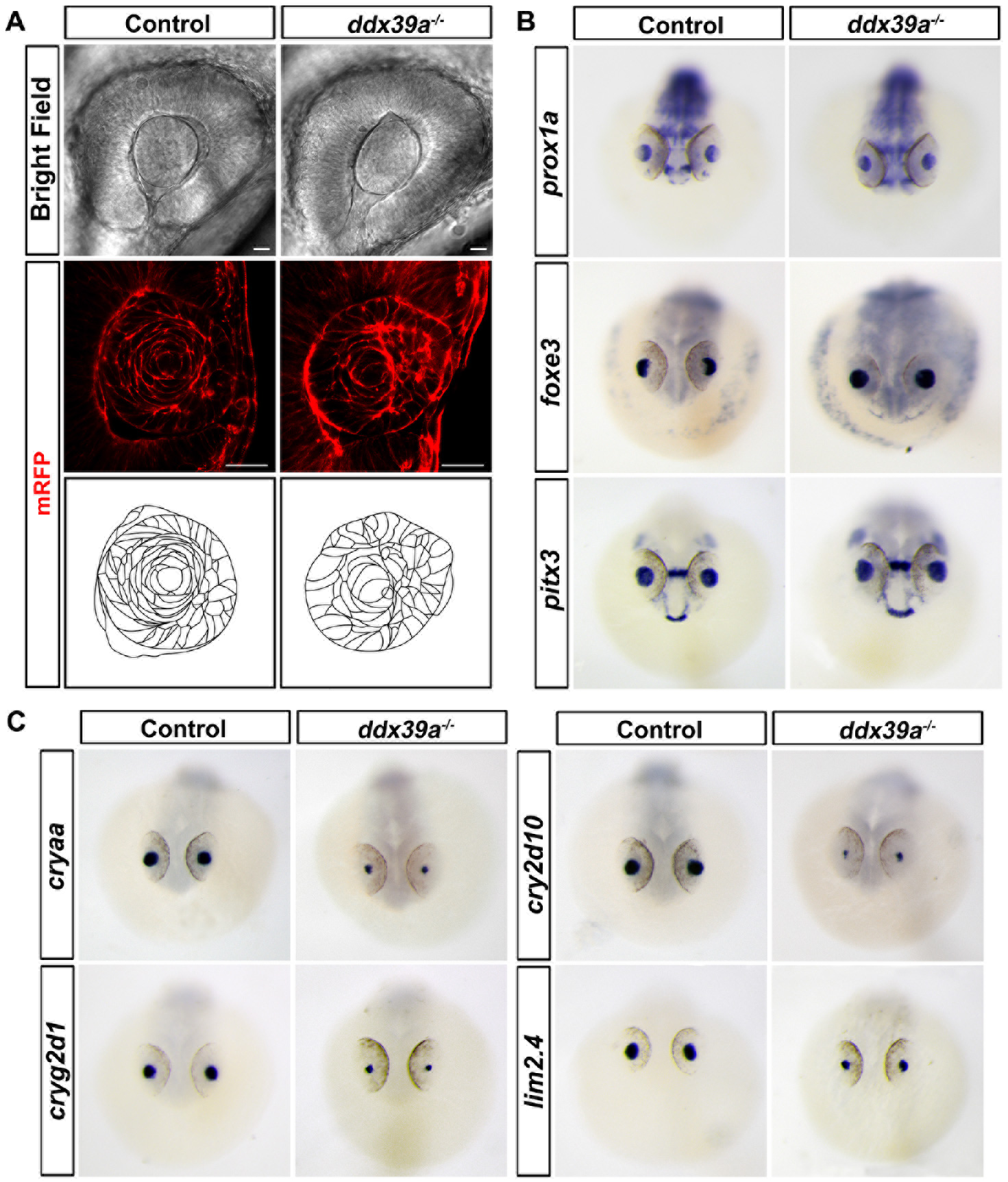
Lens fiber cell differentiation was interrupted in *ddx39a* mutant embryo. (A) : *Ddx39a* mutants possessed defects in lens fiber morphogenesis. Upper panels, bright field images of the eyes from wide type or *ddx39a* mutant zebrafish embryos at 24 hpf. Middle panels, transverse sections showed organization in lens fibers at 24 hpf, cell membrane was labeled with mRFP through mRNA microinjection. Button panels, drawings schematically show the shape and arrangement of lens fiber cells. Scale bars, 50uM. (B and C): RNA *in situ* hybridization is shown for lens gene expression in wild type and *ddx39a* mutant zebrafish embryos for indicated transcripts at 32 hpf. B and C, frontal views with dorsal side to the top.

The above results indicated ddx39a deletion causes disaregulated expression of structural genes during terminal differentiation of cardiomyocyte, myocyte and lens fiber cells. This hampered cell differentiation in turn leads to multiple defects in the muscular organs and eyes, which could collectively contribute to the lethal phenotype in *ddx39a* mutant zebrafish embryos.

### Changes in the transcriptomic landscape in ddx39a mutants

Dead box RNA helicases have been shown to exert their biological function through regulating multiple facets of RNA metabolism (Linder and Jankowsky, 2011). In order to determine how mutation of *ddx39a* affects the zebrafish embryo transcriptome, we pursued RNA-Seq on wild type and *ddx39a* mutant embryos at 24 hpf. A total of 878 genes, consisting of 548 with decreased and 330 with increased expression, were significantly altered in the *ddx39a* mutant embryo (fold change (FC) >2, FDR < 0.05) (Fig. 5A; Table S3). Following gene ontology (GO) analysis, we found that genes down-regulated in *ddx39a* mutants were enriched for GO terms linked to development of muscular tissue and lens (Fig. 5B). Interestingly, consistent with previous observations revealed by *in situ* hybridization, in all three type cells (cardiomyocyte, myocyte and lens fiber cells), the expression of upstream regulatory factors showed no evident change while the expression of many structural constituents was greatly diminished (highlighted in Fig. 5B). These results further suggested that *ddx39a* plays a common and important role in the terminal differentiation of cardiomyocyte, myocyte and lens fiber cells.

To gain further insight into the impact on transcript levels caused by loss of of *ddx39a*, we examined alternative splicing (AS) by using the OLego program and the Quantas pipeline. 12,236 significant AS events (|dI| > 0.1; Table S4) were scored in *ddx39a* mutant embryos compared with controls. These events include skipping/inclusion of single cassette exons or tandem cassette exons, intron retention and the use of alternative 5′ or 3 ′ splice sites (Table S4, number of events shown in Fig. 5C). These data showed that loss of Ddx39a function leads to extensive intron retention events and cassette skipping, implicating Ddx39a functions in intron definition. The mixed splicing activities (skipping and inclusion) observed in *ddx39a* mutants clearly suggest context-dependent splicing defects result from the loss of *ddx39a*.

**Figure 5.**
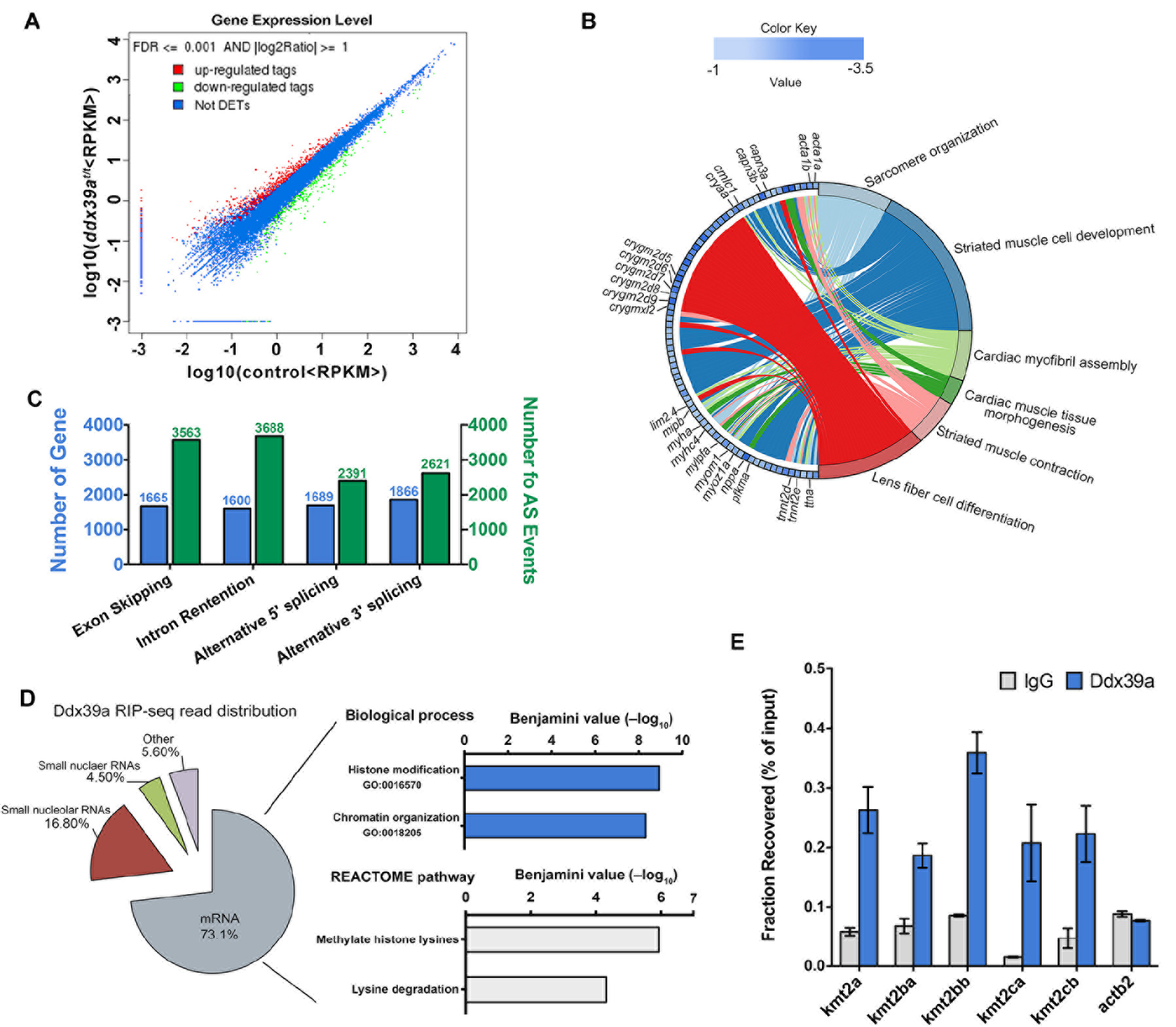
Transcriptomic landscape of *ddx39a* mutants and identification of Ddx39a-associating RNAs. (A): Expression levels of differentially expressed genes in control and *ddx39a* mutant embryo. Up-regulated and down-regulated genes are denoted by red and green spots respectively, while not differentially expressed genes are denoted as blue spots. RPKM, reads per kilobase per million mapped reads. (B): Circular plot showing 77 representative down regulated genes belonging to enriched functional categories, simultaneously presenting a detailed view of the relationships between expression changes (left semicircle perimeter) and Gene Ontology (right semicircle perimeter). Complete gene lists and GO terms tables are available in the Table S4) (C): Distribution of various alternative splicing events in *ddx39a* mutant embryos. Details in Table S5. (D): Left, Ddx39a RIP-Seq reads annotated to zebrafish genome with percentage of the total RIP-seq reads shown. Right, enriched Gene Ontology and REACTOME pathway terms from Ddx39a-bound mRNAs obtained using the DAVID tool. The x axis values correspond to the negative Benjamini P value. (E): RIP-qPCR showing binding of Ddx39a to selected epigenetic modulator genes. *Actb2* was included as negative control.

### Ddx39a deletion affects pre-mRNA splicing of KMT2 family genes

We proposed that defining the RNA interactome of Ddx39a would reveal insights into the molecular mechanisms underlying the phenotypes in *ddx39a* mutant embryos. To systematically identify Ddx39a-associated RNAs, RNA Immunoprecipitation sequencing (RIP-seq) was performed in *ddx39a* mutant embryos injected with mRNA coding Flag-tagged Ddx39a (Flag-Ddx39a). Sequencing results showed Ddx39a interacts with a diverse set of RNAs (Table S5), of which mRNAs were highly represented (73.1%), with snoRNAs and rRNA contributed 16.8% and 4.5%, respectively, of bound transcripts (Fig. 5D). Comparison of data from RIP-seq and RNA seq revealed that 84% Ddx39a-associated mRNAs showed alternative splicing in *ddx39a* mutants, underscoring the function of Ddx39a in pre-mRNA splicing. Gene ontology term and pathway analysis linked these Ddx39a-associating mRNAs to histone modification. Among these potential Ddx39a target mRNAs, we noticed members of KMT2 gene family (*kmt2a*, *kmt2ba*, *kmt2bb* and *kmt2cb*) were prominent. KMT2 family members methylate lysine 4 on the histone H3 tail, a critical regulatory step associated with myogenesis through modulation of chromatin structure and DNA accessibility (Lee et al., 2013; Rao and Dou, 2015).

To confirmed interactions between Ddx39a and KMT2 family members mRNAs, RNA immunoprecipitation and quantitative reverse transcription PCR (RIP-qPCR) was applied. This further verified that Ddx39a could bind to these mRNAs (Fig. 5E). We further examined splicing events of KMT2 family genes in control and *ddx39a* mutant embryos by using RT-PCR analysis (Rios et al., 2011; Rosel et al., 2011). This analysis showed that unspliced mRNAs were retained at higher levels in *ddx39a* mutants at 24 hpf (Fig. 6A), suggesting that the pre-mRNA splicing of these genes was defective. As a control, we confirmed that in *ddx39a* mutants the splicing of the housekeeping gene *actb1* was normal. These results suggest that in *ddx39a* mutants the effect of pre-mRNA splicing may be specific to a certain set of genes.

KMT2 enzymes are the major histone methyltransferases responsible for mono-methylation at lysine 4 of Histone 3 (H3K4me1) at distal enhancers and regions flanking the transcription start site (TSS) (Rao and Dou, 2015). To investigate whether loss of *ddx39a* affects the presence of H3K4me1 on actively transcribed genes, CHIP-qPCR was performed for a selected set of genes that showed altered transcript levels in control and *ddx39a* mutant embryos at 24 hpf. Whereas global H3K4me1 levels in *ddx39a* mutants showed only minor decreases as compared to wild type (Fig. S3), the H3K4me1 occupancy on regions flanking the TSS of myocyte and cardiomyocyte specific genes (*acta1b*, *myhz2*, *nppa*, *myom1a*, *tnnt2d*, *mylpfb* and *smhyc1*) was significantly reduced in mutants (Fig. 6B). Interestingly, the H3K4me1 occupancy level in the TSS region of myogenic regulatory factors (such as *myod*) were comparable to control embryo in *ddx39a* mutants, consistent with analysis of transcript and protein levels via RNA-seq, *in situ* hybridization and Western blot (Fig. 6B and Fig. S3).

**Figure 6.**
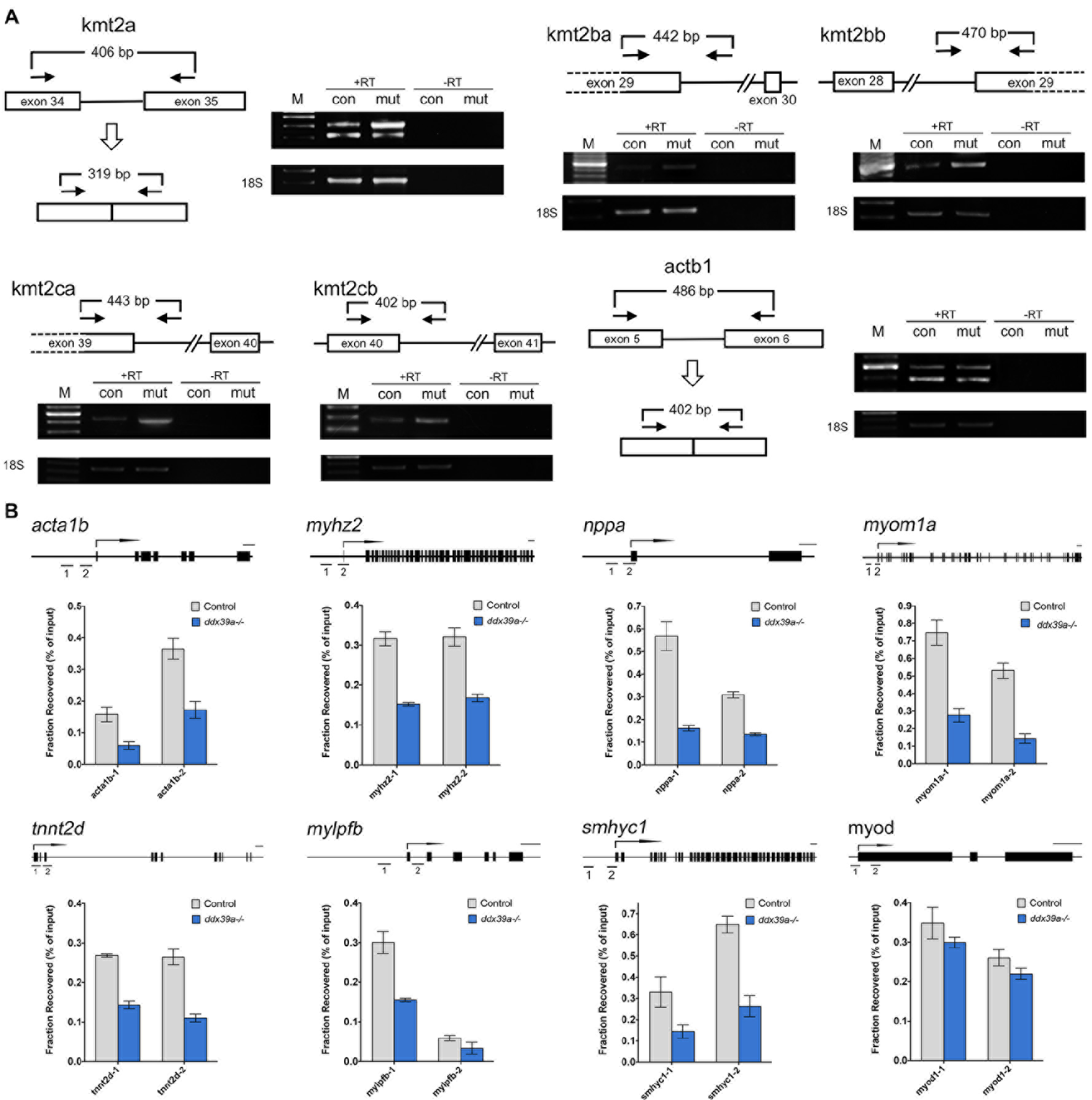
*Ddx39* deficiency affects pre-mRNA splicing of epigenetic modulator genes. (A): The splicing status of *kmt2a, kmt2ba, kmt2bb, kmt2ca, kmt2cb* and *actb1* pre-mRNA was monitored using RT-PCR with the primers indicated in schemes (boxes, exons; lines, introns; arrows, primers). Unspliced *kmt2a, kmt2ba, kmt2bb, kmt2ca and kmt2cb* mRNAs were retained in the *ddx39a* mutant (mut) larvae compared to the control (con) larvae. Unspliced and spliced PCR products were verified by sequencing. +RT refers to the validation reaction itself, and -RT represents the respective control reaction without reverse transcriptase. 18S is a loading control. M, DNA size markers.Control larvae were sibling wild type or *ddx39a*^+/−^ larvae and had normal phenotypes.(B) : ChIP-qPCR analyses represent the effect from loss of *Ddx39* on H3K4 Methylation levels at promoter and TSS of muscle cell genes. Upper schematics showed the gene structure (black rectangles, exons; arrow, TSS) and indicated the DNA segments amplified by PCR.

These results support a context-dependent requirement of Ddx39a for proper pre-mRNA splicing of KMT2 family members, with loss of *ddx39a* leading to a failure in establishment of the epigenetic status required for terminal differentiation.

## Discussion

Members of DEAD box RNA helicase family have been shown to be involved in nearly all aspects of RNA metabolism, from transcription to mRNA decay. Beyond the scale of biochemistry and cell biology, recent studies have started to delineate different functions of DEAD box RNA helicase in variant biological scenarios including animal development. For example, *ddx46* has been shown to be expressed in the digestive organs and brain and required for development of these organs (Hozumi et al., 2012). Researchers have also reported that *ddx18* is essential for hematopoiesis (Payne et al., 2011). The generation of the *ddx39a* gene-trapping allele provided the opportunity to study the function of this gene in vertebrate development. Here we demonstrated that *ddx39a* is required for normal gene expression and differentiation of cardiomyocyte, myocyte and lens fiber cells. In previous studies, development and function of heart has proven to be sensitive to defects in RNA metabolism (Ding et al., 2004; Xu et al., 2005). Our observations corroborate these results and suggest that myocyte and lens fiber differentiation similarly bear this susceptibility.

RNA-binding proteins (RBP) including DEAD box RNA helicases modulate splicing primarily by positively or negatively regulating splice-site recognition by the spliceosome. The recognition of RBP on pre-mRNAs relies on the distinct regulatory sequences in pre-mRNA that function as splicing enhancers or silencers. Our data showed Ddx39a could bind to a battery of mRNAs while more sophisticated molecular biology and bioinformatics efforts need to be exploited to unfold the detail mechanism of Ddx39a-mediating splicing events such as how is the specificity defined.

During the development, *ddx39a* shows a complex and dynamic expression pattern(Fig. 1 and Fig. S2). This result prompts future analysis of the role of *ddx39a* at later developmental events (e.g. formation of pharyngeal arches and development of digestive organs). However, the severe defects observed in *ddx39a* mutants at early stages of development complicates the analysis of later developmental events. CRIPSR/Cas9-based generation of tissue-specific *ddx39a* mutants, or generation of a conditional *ddx39a* allele, will be required to study *ddx39a* function in other organs/tissues.

Previous studies showed *ddx39a* acts as a growth-associated factor in cancer cells that is required for genome integrity and telomere protection (Sugiura et al., 2007; Yoo and Chung, 2011). We did not detect abnormal mitoses in *ddx39a* mutant embryos (Sup Fig 3). However, analysis of later stage *ddx39a* mutants may be required to observe telomeric defects, after a sufficient number of cell divisions have occurred. It also could be true that the telomere protection function of *ddx39a* is not conserved between fish and mammals.

It’s interesting to note that among a relative small number of genes significantly down-regulated in *ddx39a* mutant embryos, a large proportion encoded structural constituents including sarcomeric components in cardiomyocytes and crystallin genes in lens fiber cells. In addition to regulating mRNA splicing, it is possible that Ddx39a is involved in transactivation of structural components of via uncharacterized mechanism that is exploited in both muscular and lens fiber cells. Several studies have shown that some DEAD box proteins play important roles as regulators of transcription, particularly as co-activators or co-suppressors of transcription factors (Fuller-Pace and Nicol, 2012; Huang et al., 2015). Further investigation to determine proteins Ddx39a directly binds in various cell types, and the functional consequences of these interactions, should provide important insight into how the specificity of Ddx39a function is regulated.

## Acknowledgement

We thank Qi Xiao for drawing schematic pictures, Peipei Yin for zebrafish husbandry and members of the Scott lab at University of Toronto and Di Chen lab at Nanjing University for feedback and help during this project. This research was undertaken, in part, thanks to grant funding from National Natural Science Foundation of China (NSFC 31671505 and NSFC 31471354 to X.L), and grant funding from the Natural Sciences and Engineering Research Council of Canada (RGPIN 2017-06502 to I.C.S.).

## Author contributions

X.L. and I.C.S. conceived and supervised the project and wrote the manuscript. L. Z., B. L., Y.Y. and X.L. performed all experiments.

---

**Supplemental Figure 1.**
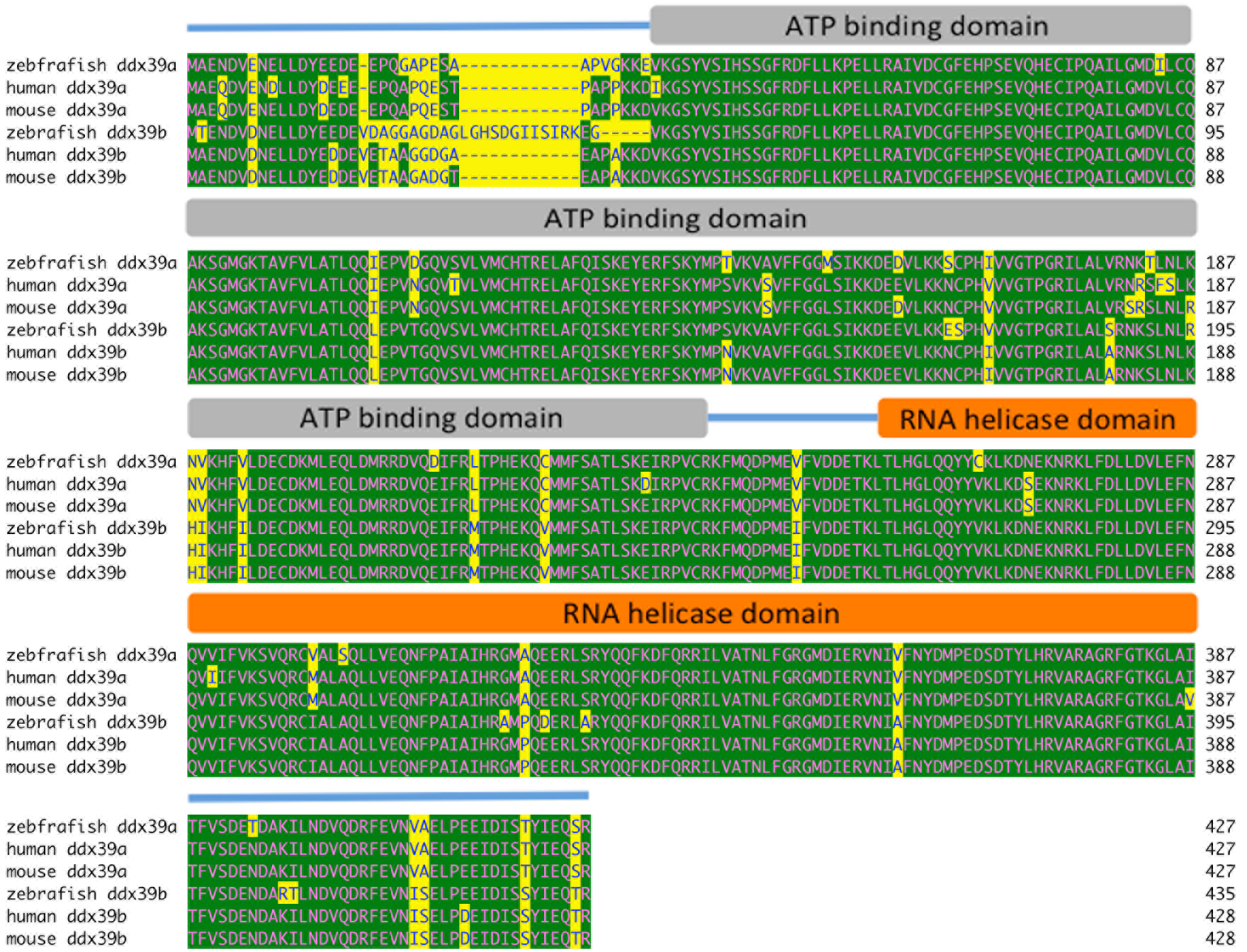
DDX39 protein sequences alignment. Functional domains are labeled. The orthologs DDX39A share 94% amino acid identity, differing in only 21 residues.

**Supplemental Figure 2.**
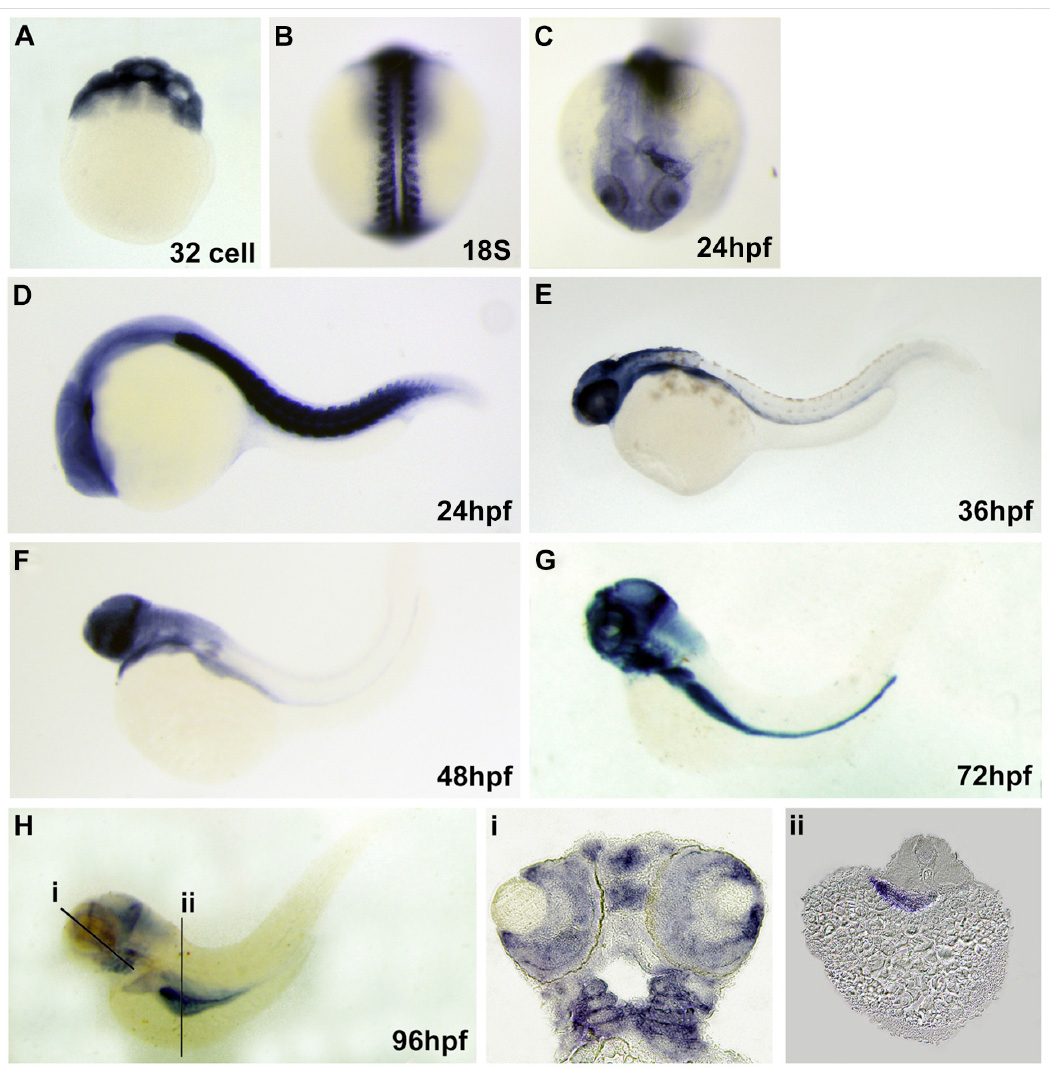
Zebrafish *ddx39a* embryonic expression pattern. (A): In situ hybridization experiment revealed there is maternal deposit ddx39a mRNA in early stage embryo. (B): Strong expression of *ddx39a* could be observed in myotome at 18 somite stage. (C-D): At 24hpf enriched *ddx39a* mRNA could be observed in lens, heart tube and trunk muscle. (E-H): Later on, the expression was restrained to specific regions in brain, retina (i), pharyngeal arches (i) and endoderm derived organs (ii). A, lateral view. B, dorsal view. C, frontal view. D to H, head to left. i and ii are sections of H as indicated. hpf, hour past fertilization.

**Supplemental Figure 3.**
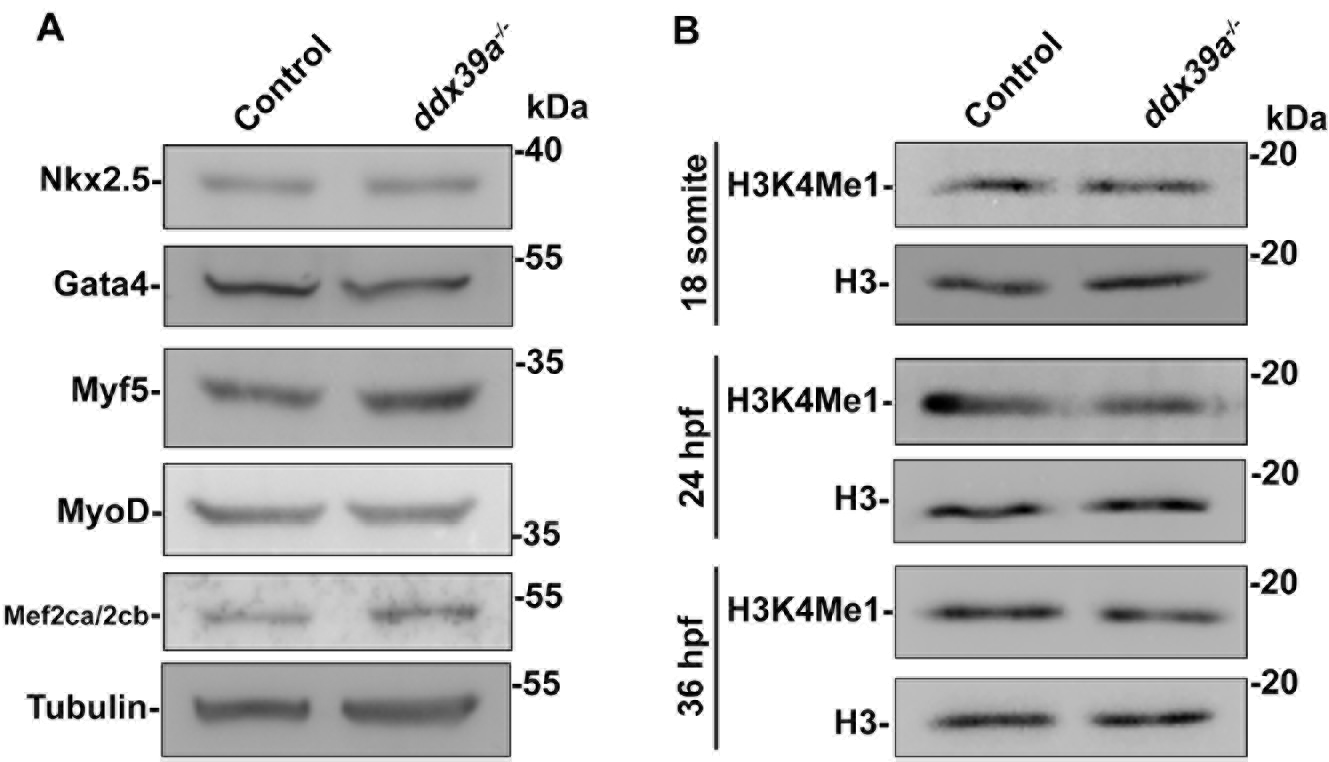
Western blot analysis of the indicated proteins and histone PTMs in control and *ddx39a* mutant embryo. (A): protein level of myocyte or cardiomyocyte key transcription regulators showed no evident change between control and *ddx39a* mutant at 24 hpf. (B): In *ddx39a* mutant, minor decrease on H3K4Me1 level could be detected from 24 hpf.

**Supplemental Figure 4.**
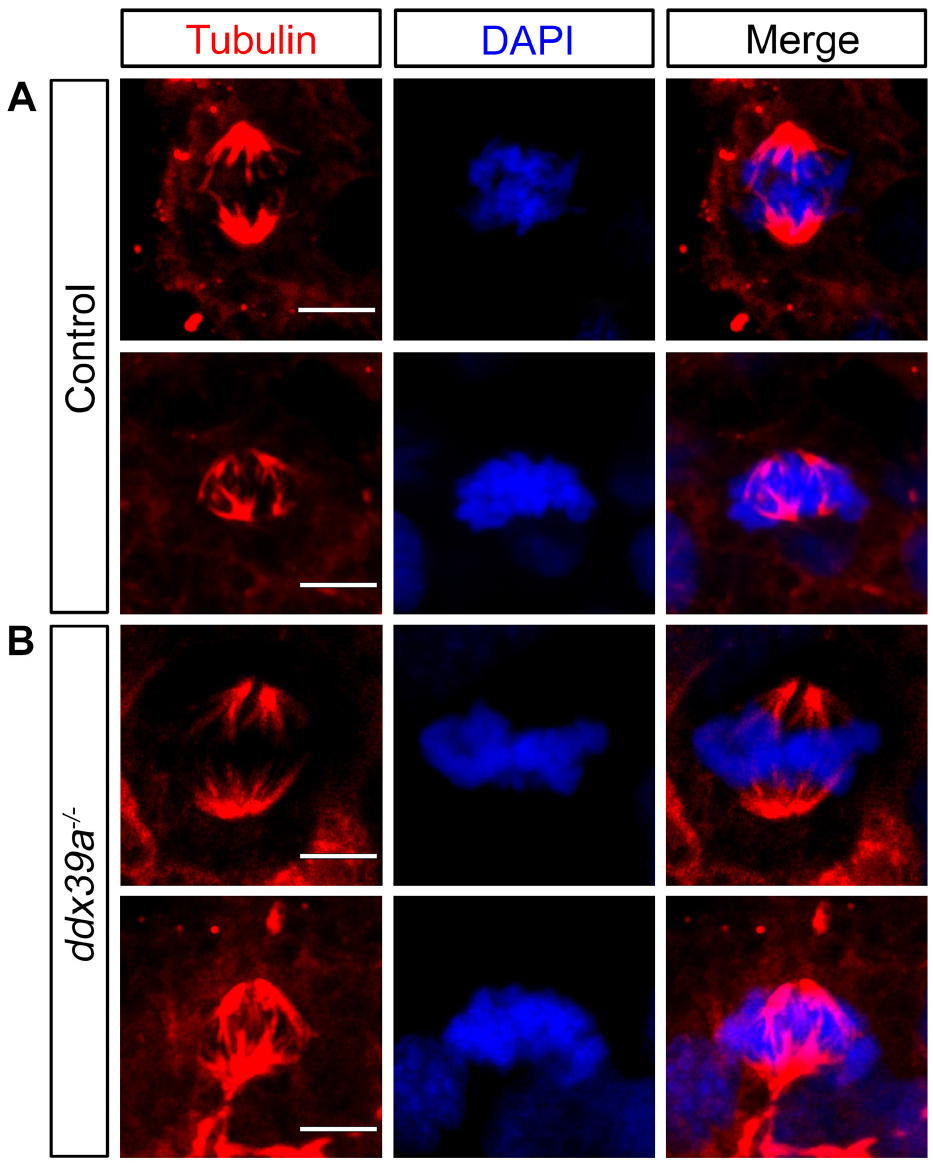
*Ddx39a* mutant embryos exhibit normal mitoses. Mitotic cells stained with DAPI (blue) and α-tubulin (red). Scale bars, 5um.

